# Integrated Computing and Tracking System for Centralized High-Throughput Genetic Analysis: a Case Study

**DOI:** 10.1101/137596

**Authors:** Adrienne M. Stilp, Stephanie M. Gogarten, Cathy C. Laurie, Tamar Sofer

## Abstract

The Genetic Analysis Center (GAC) of the Hispanic Community Health Study/Study of Latinos (HCHS/SOL) developed an Integrated Computing and Tracking system (ICT) in order to perform genome-wide and other genetic association studies automatically and efficiently, while documenting all analysis specifications. This system provides easy-to-use analysis set-up and computing procedures, automatic reports, and analysis search functionality due to integration with an on-site database. In this paper we describe the ICT and demonstrate how it satisfies key principles of reproducible research, while respecting constraints and challenges arising from using very large, restricted access, human-subjects data. This case study may benefit other groups that have similar requirements for high-throughput analysis execution and management.

## Introduction

The Genetic Analysis Center of the Hispanics Community Health Study/Study of Latinos (HCHS/SOL GAC, GAC henceforth), was established to perform genetic association analyses in collaboration with thirteen working groups, each focusing on a specific family of health outcomes and their genetic bases. To better perform and track these analyses, the GAC developed an Integrated Computing and Tracking (ICT) system for analysis execution and management to efficiently perform standardized analysis tasks, while implementing many key principles of reproducible research.

To support investigators studying the associations of genetic variants with a trait, the GAC performed many Genome-Wide Association Studies (GWAS), as well as other follow-up analyses such as generation of LocusZoom plots (Pruim et al., 2010), heritability analyses, and others (Dunn et al., 2016; Schick et al., 2016; Brown et al., 2017; Sanders et al., 2017). All primary analyses belong to a relatively small set of GWAS types: linear models, linear mixed models, logistic mixed models, etc. A unique challenge of the GAC was to efficiently perform hundreds of analyses centrally for a large group of collaborators, while ensuring standardization, reproducibility and proper methods documentation for publication. While there are existing analysis pipelines for biological data analysis (Sloggett et al., 2013; Sadedin et al., 2012; Goecks et al., 2010), they do not generally address the use of very large data sets containing protected human-subjects data. To ensure efficiency and reproducibility of research results, the GAC integrated its analysis pipeline with an on-site SQL database that tracks the input genetic data files, the phenotype data, software versions and the parameters of all analyses run by GAC analysts.

In this manuscript we describe the ICT, composed of (1) an analysis pipeline implemented using both Python scripts to manage jobs on a compute cluster and R scripts for statistical analysis; (2) an SQL database for managing genotypes, phenotypes and analysis parameters; and (3) a dedicated R package for interaction between the database and the analysis pipeline. We note that this system provides reproducible analyses in the sense that internal GAC users can fully reproduce an analysis based on information stored in the database (input parameters and software versions). This case study presents a productive implementation of many principles of reproducibility, given limitations on data sharing as well as large datasets required for genomic analyses, and may be useful to other groups with similar requirements for efficient analysis performance, tracking and management.

## Methods

### Fundamental requirements from the ICT

Based on the genetic analysis workflows, these are the core functions we required from the ICT:

#### 1. Efficient specification of a core set of analyses by the analysts

Analyses consist of a primary GWAS which include association tests and summary figures such as quantile-quantile and Manhattan plots, as well as optional follow-up analyses, such as LocusZoom plots, forest plots, and others. The primary GWAS analysis requires specification of sample set, outcome, covariates, data transformations, covariance matrices for a mixed model analysis, and the appropriate set of principal components to adjust for population stratification. Follow-up analyses use results from a specific, existing GWAS, and typically require information such as unique SNP identifiers or kinship coefficient thresholds.

#### 2. Tracking of both GWAS properties and results

For communication of scientific results, reproducibility purposes, and potential replication in other studies, it is important to track the specific variables (including transformations) and participants analyzed, inclusion and exclusion criteria leading to the specific sample of participants, and other model specifications. Tracking of analyses also allows for communication with investigators about the specific results presented in a manuscript, and for easy verifications and spot-checks if needed. It is also necessary to store the version of code used to run each GWAS for reproducibility purposes, as results could potentially depend on which software version was used for calculations.

#### 3. Searching and management of tracked analyses

The ability to work with analysis parameters in a centralized system offers useful functionality given the large number of manuscripts supported by the GAC. First, analyses can be more easily compared to each other when needed. For example, in early stages of the research, we studied which set of principal components best controlled population stratification (Conomos et al., 2016a). Therefore, we sought to compare summary measures across analyses that used the different sets of PCs. The ICT facilitated this comparison by allowing us to more easily find and process results from the two sets of analyses. Second, search capabilities are useful for fixing errors, when needed in multiple analyses. For instance, after running multiple analyses of binary outcomes using linear mixed models (LMMs), we developed a method to fit a generalized linear model (GLM) for the complex HCHS/SOL data set (Chen et al., 2016). Using search capabilities, we were able to find analyses that used LMM and re-run them using the new method. Moreover, if bugs are fixed in the analysis code, we can quickly identify and rerun analyses that used affected code versions.

#### 4. Automatic packaging and generation of reports

After running an analysis, GAC analysts provide the GWAS results to the working group investigators who requested it. The ICT allows GAC analysts to create consistent results packages with all necessary files, including data dictionaries and annotations, for any analysis with a single command. Attached reports provide both the GWAS properties as well as data provenance for investigators.

#### 5. Quality control

We sought to implement both automatic and manual procedures to check and report errors in submitted analyses. Automatic procedures in the analysis scripts can identify known problems, such as lack of convergence of a mixed model algorithm, and manual procedures can be used to perform searches once a new, previously unknown, problem arises. For instance, a few months after the GAC changed the default parameters of a GWAS to allow for heterogeneous variances among the HCHS/SOL ethnic groups, one analyst reported that a sex-stratified analysis failed. In a manual search in the database according to analyses characteristics, we found that the meta-analysis phase of a sex-stratified analysis fails when heterogeneous variances are allowed and we adapted our code accordingly. We also searched for other analyses with similar characteristics to report this problem to the analysts who ran them.

### The ICT design

There are two major components to the ICT. The first is a database that holds information on all data components (traits, principal components, location of genotype data files, etc.), and on all specific analyses that are run. The second component is code that interacts with the database to set up the analyses, run both the GWAS and follow up analyses, package the results, and provide an interface to the analysis results for interactive work by GAC analysts.

Figure 1 portrays a workflow of an analysis and its interaction with the ICT. An analysis starts with a preparation stage, in which traits are selected, exclusion criteria are applied, model parameters are tuned, and checks are performed for model fit. The analyst prepares a set of configuration files in which all information necessary for the analysis is specified. Next an analysis shell is created with a standard directory structure and information about the analysis is written in the database, including the assignment of a unique analysis id that is used for all future interactions with this analysis. Finally, the analysis is run and results, as well as figures and a report, are written to files in the previously built directories. Additional follow-up analyses may be run later.

**Figure 1:**
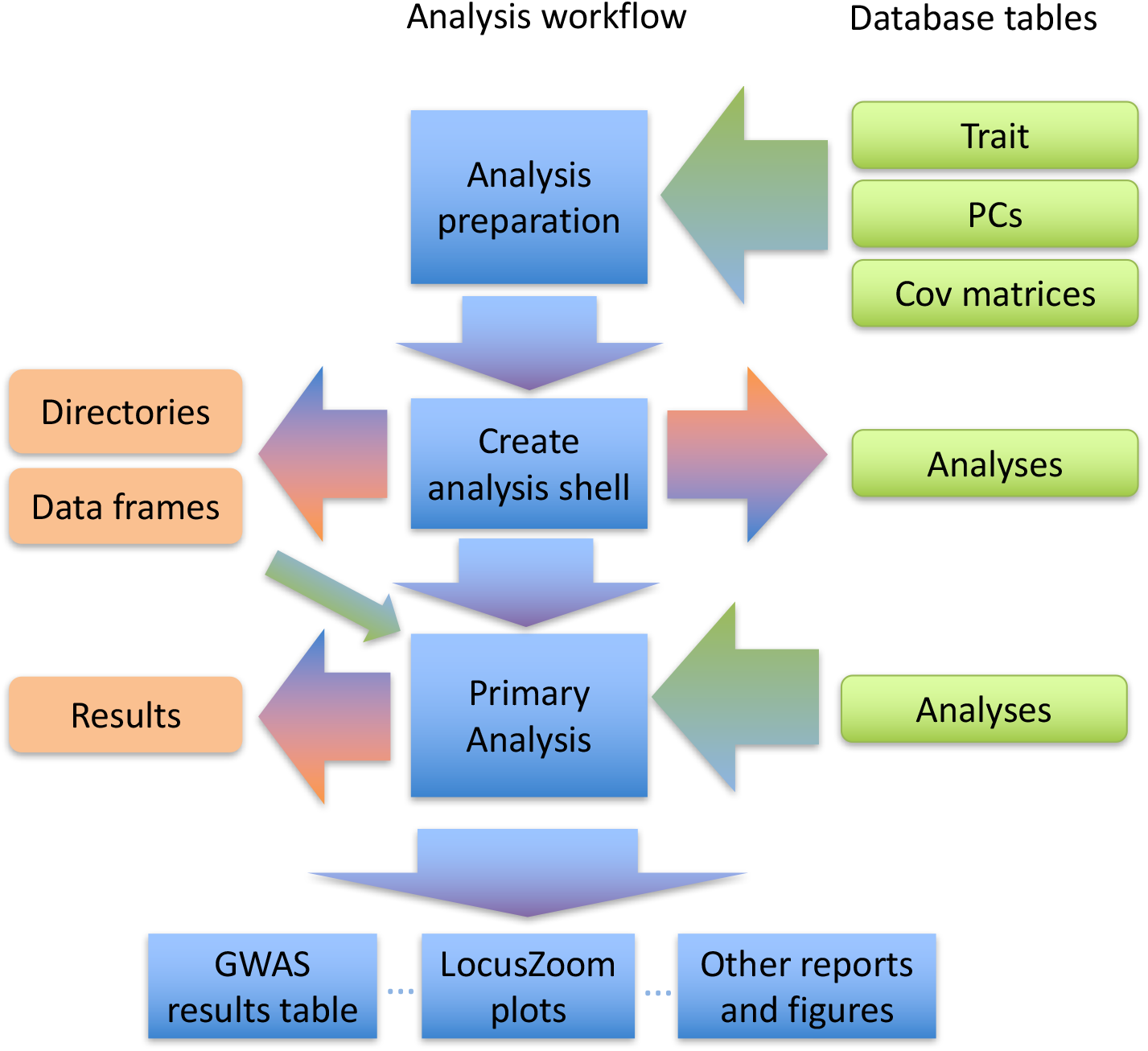
A diagram of a workflow of an analysis, and the interaction of the analysis with the database. Incoming arrows to the middle column indicates data read from the database, and outgoing arrows indicates writing of information to the database and to the file system.

#### The database

Table 1 provides a list of database tables and a brief description of their function. The database schema is provided as S1 Figure in the supporting information. The database stores all information necessary to recreate an analysis. It also stores phenotype data and the principal component sets describing population structure, with unique identifiers assigned by the database. Larger data, such as covariance matrices or genotype datasets, are not stored directly in the database but instead are identified by a path to their location on disk. Unique identifiers for each matrix or genotype dataset, and for their various versions, are assigned. GAC analysts use these unique identifiers (e.g., trait id, principal component id, matrix id) to pull data from the database and to prepare setup files for an analysis. The database also stores the exact sample set used in the analysis after excluding participants for reasons relating to the phenotype being tested (e.g., extreme phenotype values or medication use) or because they had missing data in any of the covariate models. When an analysis and its details are added to the database, a unique analysis id is automatically assigned to that analysis, which is then used by all analysis scripts, and by all functions querying the database regarding the analysis and its results. Finally, the version of the code that is used for an analysis is stored in the ICT analysis table.

**Table 1:**
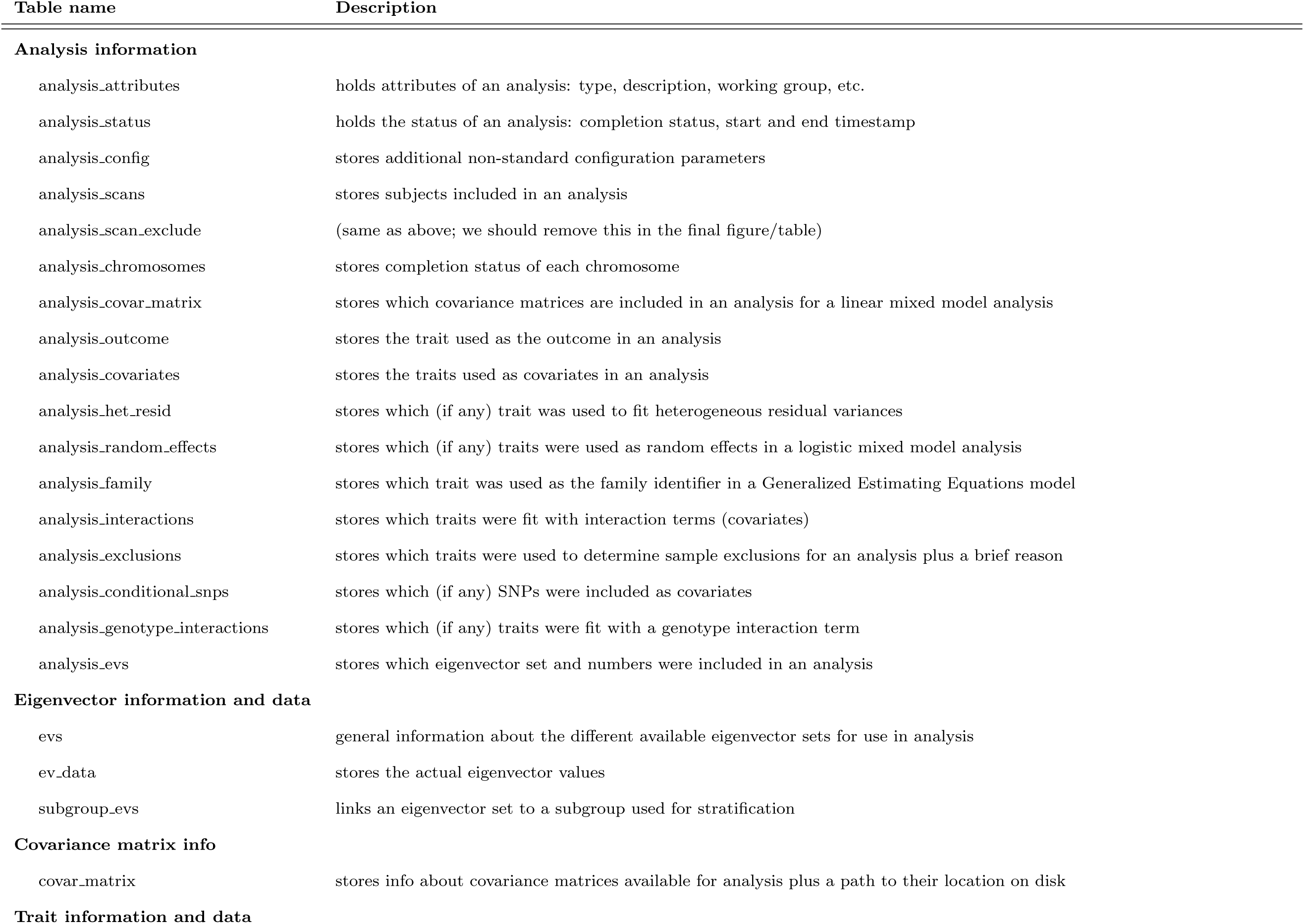
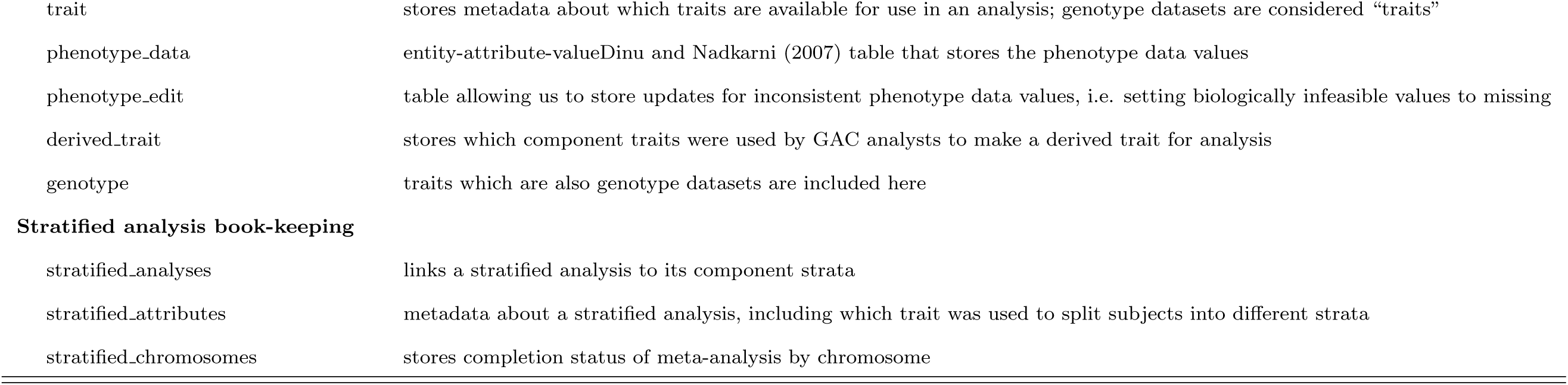
Description of database tables

| Table name | Description |
| --- | --- |
| **Analysis information** |  |
| analysis_attributes | holds attributes of an analysis: type, description, working group, etc. |
| analysis_status | holds the status of an analysis: completion status, start and end timestamp |
| analysis_config | stores additional non-standard configuration parameters |
| analysis_scans | stores subjects included in an analysis |
| analysis_scan_exclude | (same as above; we should remove this in the final figure/table) |
| analysis_chromosomes | stores completion status of each chromosome |
| analysis_covar_matrix | stores which covariance matrices are included in an analysis for a linear mixed model analysis |
| analysis_outcome | stores the trait used as the outcome in an analysis |
| analysis_covariates | stores the traits used as covariates in an analysis |
| analysis_het_resid | stores which (if any) trait was used to fit heterogeneous residual variances |
| analysis_random_effects | stores which (if any) traits were used as random effects in a logistic mixed model analysis |
| analysis_family | stores which trait was used as the family identifier in a Generalized Estimating Equations model |
| analysis_interactions | stores which traits were fit with interaction terms (covariates) |
| analysis_exclusions | stores which traits were used to determine sample exclusions for an analysis plus a brief reason |
| analysis_conditional_snps | stores which (if any) SNPs were included as covariates |
| analysis_genotype_interactions | stores which (if any) traits were fit with a genotype interaction term |
| analysis_evs | stores which eigenvector set and numbers were included in an analysis |
| **Eigenvector information and data** |  |
| evs | general information about the different available eigenvector sets for use in analysis |
| ev_data | stores the actual eigenvector values |
| subgroup_evs | links an eigenvector set to a subgroup used for stratification |
| **Covariance matrix info** |  |
| covar_matrix | stores info about covariance matrices available for analysis plus a path to their location on disk |
| **Trait information and data** |  |
| trait | stores metadata about which traits are available for use in an analysis; genotype datasets are considered “traits” |
| phenotype_data | entity-attribute-valueDinu and Nadkarni (2007) table that stores the phenotype data values |
| phenotype_edit | table allowing us to store updates for inconsistent phenotype data values, i.e. setting biologically infeasible values to missing |
| derived_trait | stores which component traits were used by GAC analysts to make a derived trait for analysis |
| genotype | traits which are also genotype datasets are included here |
| **Stratified analysis book-keeping** |  |
| stratified_analyses | links a stratified analysis to its component strata |
| stratified_attributes | metadata about a stratified analysis, including which trait was used to split subjects into different strata |
| stratified_chromosomes | stores completion status of meta-analysis by chromosome |

#### Analysis submission scripts and code

We developed an analysis R package containing statistical analysis software (the mixed model code from this package was later incorporated into the GENESIS (Conomos et al., 2016b) and MetaCor (Sofer et al., 2016) packages), and a second R package to interact with the database by recording analysis details. extracting data values, and reading analysis results. Because we are using protected human-subjects data in our analyses, the database is only accessible from GAC computers. The analysis code also includes scripts that call functions within the R packages to obtain analysis configuration information, run analyses, and write files to disk.

Before running an analysis, the analysts first runs a script in the ICT that checks setup files for consistency, and generates diagnostic plots to assess the quality of a basic model that includes all adjusting covariates and random effects (if needed), but excludes the genetic variants that will be tested later. The analyst inspects the diagnostics tests and verifies that there are no errors, e.g. a singular design matrix, and then directs the ICT, by running a script, to add the analysis to the database. In turn, the ICT adds appropriate entries to the database and builds directories. Once the analysis is added, it is identified using its automatically-assigned unique analysis id. The analysis jobs are then submitted to the cluster using Python scripts, which perform some general checks (e.g. Is the submitting user the one who owns the analysis? Have certain pieces of the analysis already been run?) and submits the appropriate R scripts, which are different for analyses using different statistical tests. For example, mixed model analyses require estimation of variance components, but linear model analyses do not. For all analyses, a report is automatically generated using a Sweave (Leisch, 2002) template that records all details of the given analysis based on the information stored in the database (see S1 File for an example report). After all standard analysis scripts have finished, a final script is run that checks for the success of all jobs, organizes the output files, updates the database with the analysis status (i.e. completed or failed), and write-protects the directory to make it more difficult for a user to accidentally delete or overwrite results. Follow-up analyses may be run via additional Python submission scripts, all of which require the unique analysis identifier of the primary analysis.

#### Reproducible analyses

The analysis report documents analysis properties and components, and is helpful in reproducing research (running the same analysis). A more precise and automated way for reproducing an analysis is via an R command that obtains all properties of a specific analysis using its analysis identifier and parameters stored in the database. This function generates all configuration files allowing for running the same analysis, which can be submitted as is or easily altered to specify a similar analysis based on the original analysis. For instance, one may need to run the same analysis but with additional sample exclusions as a sensitivity analysis or with additional adjusting variables (e.g. a genetic variant that was already associated with the outcome for a conditional association analysis).

#### Data integrity verification

In addition to checks immediately before and after an analysis is submitted, we also run a nightly cron job to check for consistency between the database and analysis results. This script verifies that all input files tracked in the database exist (e.g., genotype datasets and covariance matrix files); that all analyses in the database have a corresponding directory on disk; that all analysis directories on disk have a corresponding entry in the database; and that all chromosomes marked “completed” in the database for a given analysis have results files in the proper location on disk. This setup provides timely warning of potential problems. For example, it has occasionally uncovered unexpected, accidental changes made by an analyst, e.g., renaming an analysis directory or moving an analysis directory to a different directory. The GAC is able to quickly remedy these problems due to automated detection and alerts by the cron job.

## Discussion

In this manuscript, we describe the ICT implemented in the HCHS/SOL GAC, a center that provided large scale genetic analysis services for HCHS/SOL investigators. Our system was designed to facilitate fast, simple, high-quality, and well-documented analyses to our community of collaborators while adhering to many principles of reproducibility. While the GAC’s system is tailored to running GWAS, the principles behind the ICT can be applied to any pipeline designed to run and track sets of similar analyses.

Reproducibility is first and foremost a tool for enforcing transparency and allowing for error detection. These characteristics are built into our ICT. Many researchers promoting reproducible research have suggested complete data availability with publication of manuscripts, so that other investigators could repeat the same analysis. This standard is difficult to implement with human subject data and large, genotypic datasets, despite data availability in controlled-access repositories such as dbGaP. For example, the same dataset that was used in one analysis may not be available in exactly the same form for future investigators due to ongoing changes in study participants’ consent or updates to the data themselves. Another limitation for such “complete reproducibility” is the change in software over time, and the use of internal packages. Although in the GAC we track all versions of our data, our analysis packages are still internally implemented, and the public packages that grew from this internal code may be slightly different from our pipeline code used at any given time (even though the statistical algorithms may be the same). Therefore, our version records will not be as helpful for external investigators. This drawback could be alleviated in future work by publishing all internal analysis code in public repositories from the beginning of the project, even if it is in the research stage, but it would still not solve problems arising from the changing nature of human-subjects datasets.

Another limitation for reproducibility is that the overall process from the first analysis of association of a trait with genetic variants to publication of a manuscript is not completely in-house. For instance, after GWAS is run by the GAC, results are transferred to investigators on a different study site, who may meta-analyze the results with results from other studies, use summary statistics for additional analyses, or apply some filtering on the analysis results. This model requires the cooperation of external investigators at multiple institutions, who typically do not have a standardized system like the ICT in place, and the GAC is unable to track their post-GWAS steps in the same robust way as the original GWAS.

Over the course of the project, we have identified some characteristics that could lead to future improvements in the reproducibility and the scientific utility of the ICT. We list them here with a brief explanation and discussion for each.

- **Store GWAS results in the database.** Currently, results for an analysis are stored as chromosome-level files within a results subdirectory of the analysis directory. While the analysis results and directory are both write-protected, it is theoretically possible for these files to be corrupted or accidentally deleted. These problems are mitigated by backups and the nightly cron job that checks for the existence of the expected results files. If the results were stored in the database, the analysis results will be subject to the same conditions as the rest of the analysis meta-data, increasing the data integrity. From a scientific standpoint, it would also be easier to run complex queries about specific results, such as cross-analysis exploration. For example, upon finding a significant association in one analysis, one could quickly find any other related outcomes that also had a significant association in the same region. However, storing results in the database presents logistical challenges. The set of results stored for each genetic variant may differ depending on the type of test used (e.g., a score test versus a Wald test), so the database schema would need to be designed to accommodate some differences based the statistical test used.
- **Require investigators to provide analysis identifiers in published manuscripts.** Due to the collaborative nature of the GAC, there is a missing link in data accessibility between the database, which is accessible only by GAC analysts, and external investigators, who are often the main authors of publications. Including the analysis identifier in publications would allow better follow-up analyses, as, given the need, a GAC analysts could more immediately pull analysis details or results using the analysis identifier linked in the publication. We have done this with some, but not all, publications resulting from our analyses.
- **Build an interface allowing external investigators to access a given set of analyses.** A direct interface for the external investigators to look up both details and results for their analysis may reduce potential miscommunication errors between GAC analysts and investigators. Such an interface would require a substantial amount of human time to build, especially considering the security restrictions required for accessing the data.
- **Better job tracking and error recovery.** Currently, only one version of the code used and only some stages of the analysis are tracked in the database. Jobs can fail due to memory issues on a compute node, but recovery is difficult because only certain stages are tracked. A future improvement would be to add a table that tracks exactly which jobs were submitted, which version of the code was used for each job (if some pieces were run with updated versions), job completion status (currently running, success, warning, or failure), and the output log file. The Python script to submit analysis jobs given an analysis id could then inspect that table and submit only the jobs necessary to complete the analysis. This proposed table could also provide better tracking of followup analyses such as Locus Zoom plots or heritability analyses, which are currently not recorded in the database.

To summarize, we provide an example of the ICT that allows for efficient and automated computation of large scale association testing, recording of the analyses results, and generation of reports. In addition to increasing the efficiency of the analyses performed in the Genetic Analysis Center, the ICT also increases the integrity of the data due to the link between the analysis pipeline and the trait and analyses database, and the ability to search for analysis properties. As we expect that such systems will be more and more common due to the increasingly large modern data sets, our case study can benefit others in the design of their systems.

## Supporting information

**S1 Fig. Database schema.**

**S1 File. Example analysis report.**

## Acknowledgments

The HCHS/SOL Genetic Analysis Center was supported by NHLBI contract HHSN268201300005C.

